# Loss of genetic variation and sex determination system in North American northern pike characterized by whole-genome resequencing

**DOI:** 10.1101/2020.06.18.157701

**Authors:** Hollie A. Johnson, Eric B. Rondeau, Ben J. G. Sutherland, David R. Minkley, Jong S. Leong, Joanne Whitehead, Cody A. Despins, Brent E. Gowen, Brian J. Collyard, Christopher M. Whipps, John M. Farrell, Ben F. Koop

**Affiliations:** Department of Biology, Centre for Biomedical Research, University of Victoria, Victoria, British Columbia, V8W 3N5, Canada; Sutherland Bioinformatics, Lantzville, British Columbia, Canada V0R 2H0; Alaska Department of Fish and Game, Division of Sport Fish, 1300 College Rd, Fairbanks, Alaska, 99701-1599, USA; Center for Applied Microbiology, Department of Environmental Biology, SUNY College of Environmental Science and Forestry, Syracuse, New York, 13210, USA; Thousand Island Biological Station, Department of Environmental and Forest Biology, SUNY College of Environmental Science and Forestry, Syracuse, New York, 13210, USA

**Keywords:** *Esox lucius*, genetic variation, genomics, long-read assembly, northern pike, population genomics, sex determination, whole-genome resequencing

## Abstract

The northern pike Esox lucius is a freshwater fish renowned for having low genetic diversity but ecological success throughout the Northern Hemisphere. Here we generate an annotated chromosome-level genome assembly of 941 Mbp in length with 25 chromosome-length scaffolds using long-reads and chromatin capture technology. We then align whole-genome resequencing data against this reference to genotype northern pike from Alaska through New Jersey (n = 47). A striking decrease in genetic diversity occurs along the sampling range, whereby samples to the west of the North American Continental Divide have substantially higher diversity than populations to the east. As an example, individuals from Interior Alaska in the west and St. Lawrence River in the east have on average 181K and 64K heterozygous SNPs per individual, respectively (i.e., a SNP variant every 3.2 kbp and 11.2 kbp, respectively). Even with such low diversity, individuals clustered with strong support within each population, and this may be related to numerous private alleles in each population. Evidence for recent population expansion was observed for a Manitoba hatchery and the St. Lawrence population (Tajima’s D = -1.07 and -1.30, respectively). Non-uniform patterns of diversity were observed across the genome, with large regions showing elevated diversity in several chromosomes, including LG24. In populations with the master sex determining gene amhby still present in the genome, amhby is in LG24. As expected, amhby was largely male-specific in Alaska and the Yukon and absent southeast to these populations, but we also document some amhby(-) males in Alaska and amhby(+) males in the Columbia River. This indicates that rather than a discrete boundary after which amhby was lost in North America, there is a patchwork of presence of this system in the western region. These results support the theory that northern pike recolonized North America from refugia in Alaska and expanded following deglaciation from west to east, with probable founder effects resulting in loss of both neutral and functional diversity including the loss of the sex determination system.

## Introduction

Historical population dynamics and contemporary genetic diversity and structure of freshwater fishes in the Northern Hemisphere have been significantly shaped by Pleistocene glacial cycles of expansion and retreat that occurred up to around 14,000 years ago (Bernatchez & Wilson, 1998; Skog et al., 2014; Wooller et al., 2015). Glacial advances resulted in large mortalities, range compressions to glacier edges, and isolation of populations into refugia. Glacial retreats resulted in formations of large postglacial lakes allowing rapid range recolonization across large geographic scales (see Bernatchez & Wilson, 1998). Northern pike *Esox lucius* (Order: Esociformes), with a distribution across much of the Northern Hemisphere in fresh and brackish water (Craig, 2008), was significantly impacted by these glacial cycles (Wooller et al., 2015). In eastern Eurasia, northern pike survived in a Siberian refugium and expanded into Beringia (Bachevskaja et al., 2019), which remained unglaciated during the Pleistocene and was a refugium for many freshwater species (reviewed by Wooller et al., 2015). Following glacial retreat, the Beringia refugium was the likely source for northern pike recolonization of Alaska (Crossman & Harington, 1970). However, Esocid ancestors of northern pike were in North America during the Paleocene (i.e., 56-66 mya) (Wilson, 1980), and therefore had a long history in the area prior to the glacial impacts (Wilson et al., 1992).

Genetic variation is considered pivotal for adaptation (Barrett & Schluter, 2008; Höglund, 2009), but northern pike throughout their range have low genetic diversity (Skov & Nilsson, 2018). Genetic diversity of northern pike is particularly low in central North America (Bosworth & Farrell, 2006; Miller & Kapuscinski, 1996; Rondeau et al., 2014; Senanan & Kapuscinski, 2000), in concordance with expectations due to glacial impacts on genomic diversity (Bernatchez & Wilson, 1998) resulting in a ‘younger’ and more recently recolonized population (reviewed by Skog et al., 2014). Although northern pike also have low diversity elsewhere in their range, including in Europe (Nicod et al., 2004), higher levels than those observed in North America occur in Sweden (Sunde et al., 2022), China (Luan et al., 2021), and Siberia (Senanan & Kapuscinski, 2000). Globally, the generally low diversity is likely due to ice age population bottlenecks and founder effects, small effective population sizes, restricted gene flow between populations, and the role of the pike as an apex predator, including with cannibalistic tendencies (Seeb et al., 1987; Skog et al., 2014). This low genetic diversity and structure challenges fine-scale population structure characterization in northern pike in North America (Miller & Senanan, 2003; Senanan & Kapuscinski, 2000; Skog et al., 2014). In general, currently three main northern pike haplogroups exist across the Northern Hemisphere (Skog et al., 2014). One of these haplogroups is Holarctic, and is present across North America as the likely expansion from an Alaskan, postglacial source described above (Wooller et al., 2015). Even with low genetic diversity, and involving only a few individuals, northern pike excel at accessing and colonizing new regions (Luan et al., 2021). Considering this historical context, and in addition to its value in sports fishing (DFO, 2012) and as a model for physiological, toxicological, and ecological studies (Forsman et al., 2015), the northern pike is a good model system to understand adaptability and resilience with low genetic diversity.

Northern pike has also been the focus of studies on mechanisms of sex determination. Pan and co-workers (2021) characterize global dynamics of the master sex determination (MSD) system in northern pike, where a male-specific duplicate of *anti-mullerian hormone*, termed *amhby*, was identified as the MSD gene in Europe, but has been lost in parts of North America. Furthermore, no replacement MSD system for *amhby*(-) populations in North America has been identified (Pan et al., 2021), which explains previous challenges in mapping sex in northern pike populations of eastern North America (see Rondeau et al., 2014). This MSD system variation is notable given that it is considered a single, circumpolar species (Grande et al., 2004). Teleosts are known to have high diversity and turnover of sex determination systems (Pan et al., 2019), and their sex determination can involve environmental factors (Devlin & Nagahama, 2002; Goto-Kazeto et al., 2006). As *amhby* is the MSD gene in Alaska, and is missing in other North America populations, this may indicate that it was lost during range expansion through bottleneck or founder effects from Alaska eastward (Pan et al., 2021). This evolving MSD system in northern pike adds to its value as a model species. Field and laboratory observations of skewed sex ratios in some North American populations do suggest an environmental effect on sex determination (Carbine, 1942; Clark, 1950; Huffman et al., 2014; Priegel & Krohn, 1975) although field-based estimates of sex ratio may contain biases (Casselman, 1975). Finer-scale resolution of populations in North America will benefit the understanding of *amhby* loss and may provide more information to understand current or evolving mechanisms underlying sex determination in *amhby*(-) populations.

Genetic resources are available for northern pike including reference genome assemblies, a linkage map, and expressed sequence tag libraries (Leong et al., 2010; Pan et al., 2019; Rondeau et al., 2014), as well as genome assemblies of other *Esox*, *Dallia*, *Novumbra*, and *Umbra* spp. (Pan et al., 2021), further adding to the value of northern pike as a model. Short-read technology has provided much insight in ecology and evolution, but long-read technology such as PacBio and Oxford Nanopore (Eid et al., 2009; Stoddart et al., 2009) can provide a more contiguous assembly that increases the potential to fully characterize genes, repeat regions, and chromosomal structural elements (reviewed by Chaisson et al., 2015). This is particularly relevant for areas of high repeat content (e.g., Bongartz, 2019) or for polyploid species in order to resolve haplotypes (e.g., Aury et al., 2022; Sun et al., 2022; Yuan et al., 2021). Long reads require error-correction with short reads (e.g., Goodwin et al., 2015; Koren et al., 2012) or consensus building with sequencing depth (Chin et al., 2013). Furthermore, new advances in scaffolding can facilitate further improvements in contiguity for super scaffolds and long-range haplotypes including chromatin conformation capture (Dekker et al., 2002) as applied in chromatin proximity ligation methods (Dudchenko et al., 2017; Mostovoy et al., 2016; Putnam et al., 2016).

In the present study we provide a new genome assembly for northern pike using long-read technology and chromatin conformation capture methods, using naming conventions consistent with the original assembly (Rondeau et al., 2014). We then use the improved assembly to characterize the genomic diversity of northern pike throughout North America using whole-genome resequencing with a particular focus on the region from Alaska through British Columbia due to its importance regarding the loss of *amhby* (Pan et al., 2021). Furthermore, we characterize the intrachromosomal variation in polymorphic and repetitive element content. Alongside this characterization of genomic variation, we use recently developed (Pan et al., 2021) and newly developed sex markers combined with histology to identify which populations have *amhby*(+) males. Collectively, this work improves our understanding of the loss of genomic variation and MSD function in a west-to-east pattern in North America, likely related to the expansion and recolonization of northern pike into North America.

## Methods

### Reference genome sampling, sequencing, and assembly

Northern pike were sampled near Castlegar, BC in September 2017 as part of an invasive species removal project in the Canadian portion of the Columbia River. Following euthanization, the liver tissue from a female individual was removed by dissection and frozen on dry ice for 48 hours before long-term storage at -80°C.

High-molecular weight DNA was extracted from the liver using a modified dialysis method as follows. Approximately 550 mg of frozen tissue was ground into a powder using liquid nitrogen and mortar and pestle. The powder was transferred to a 5 ml lo-bind tube (Eppendorf), along with 3600 μl buffer ATL, 400 μl proteinase K solution and 40 μl RNAse A solution (QIAGEN), followed by digestion at 56°C for 3 hours, with rotation at approximately 4 RPM. The homogenate was split equally into two 5 ml tubes, where a phenol-chlorform-isoamyl alcohol (25:24:1) purification was performed three times per tube, followed by a chloroform-isoamyl alcohol (24:1) purification. In each stage, one volume of organic solvent was mixed with one volume of aqueous solution, mixed by slow inversion for three minutes, then spun for 15 minutes at 5000xg for phase separation. The aqueous top layer was then transferred slowly to a new tube using a 1000 ul wide-bore pipette tip. Subsequently, 2 μl of RNAse A (20 mg/ml; QIAGEN) was added to the aqueous solution and incubated for one hour at room temperature, then 5 μl of proteinase K (20 mg/ml) was added and incubated overnight at 4°C. Finally, approximately 750 μl was obtained from each tube (i.e., 1,500 μl total) and transferred to a Spectra/Por Float-A-Lyzer G2 1000 kD (pink) dialysis device (Spectra/Por). Dialysis was performed in 1 L of 10 mM Tris-Cl (pH 8.5) at 4°C with gentle mixing for one week, changing the buffer five times throughout the process. DNA quantity was determined by Qubit v2.0 (Life Technologies) and quality by 0.6% agarose gel electrophoresis at 60 Volts. Bands greatly exceeded the largest ladder band of 40 kb with no visible shearing.

Subsequent library preparation and sequencing was performed by the McGill University and Genome Quebec Innovation Centre. In brief, PacBio sheared large-insert libraries were constructed following standard protocols and sequenced across eight SMRT cells on a PacBio Sequel, generating 76 Gbp of data. An additional library was constructed using the same input genomic DNA for 10X chromium sequencing following standard protocols and sequenced on one lane of Illumina HiSeqX PE150. A third library preparation method was conducted at the University of Victoria using the Proximo Animal Hi-C kit (Phase Genomics) following the Phase Genomics protocol 1.0 with adapter barcode N702 using 0.2 g of the liver tissue and sequenced within a single lane of Illumina HiSeq4000 PE100.

PacBio data was assembled using Canu v1.8 (Koren et al., 2017). All subreads were used, and a genome size of 0.95 Gbp was estimated as input. All stages were run with SLURM scheduling on the Compute Canada heterogeneous cluster (Cedar). Options *stageDirectory* and *gridEngineStageOption* were used for optimal on-node storage at heavy input/output stages. Default settings were used except that *ovlMerThreshold* was set to 2000, *corMhapSensitivity* was set to normal, *correctedErrorRate* to 0.085, and *minReadLength* to 2500 to reduce runtime and/or to use recommendations for Sequel data as per software guidelines. Following the initial assembly, Arrow v2.2.2 was used in SMRTlink (6.0.0.47841) to polish with PacBio data, using the ArrowGrid wrapper (Koren et al., 2017). The ArrowGrid pipeline was run a total of three times, then the output was subjected to a round of polishing using Pilon v1.22 (Walker et al., 2014) using the 10X chromium data that had been pre-processed using Longranger v2.2.2 (10X Genomics) to remove barcodes.

Scaffolding occurred in three stages. In the first stage, Hi-C data was used to scaffold the assembly with SALSA2 (Ghurye et al., 2017; Ghurye et al., 2019). Hi-C data was first prepared and aligned to the input genome following recommendations by Arima Genomics (Arima Genomics, 2023). In brief, paired-end reads from the Hi-C library were separately aligned to the genome using BWA mem 0.7.13-r1126 (Li, 2013), then sorted, merged and filtered with SAMtools v.1.8 (Li et al., 2009), Picard v.2.9.0-1-gf5b9f50 (Broad Institute, 2023), and bedtools v.2.27.0 (Quinlan & Hall, 2010). SALSA2 was then run with options *-m* yes and *-e* GATC to identify misassemblies in the input assembly and to indicate the applied restriction enzyme, respectively, and indicating the post-Pilon reference and the bed file output by the Arima pipeline.

In the second scaffolding stage, the Tigmint-ARCS-LINKS pipeline was applied. In brief, Tigmint v1.1.2 (Jackman et al., 2018) was run using the ARCS pipeline to run all three stages. The Tigmint portion of the pipeline was run with default parameters. For ARCS v1.0.5 (Yeo et al., 2017) and LINKS v1.8.6 (Warren et al., 2015), parameters were used as default except *-l* (i.e., the minimum number of links, or k-mer pairs, required to compute a scaffold; default = 5), and *-a* (i.e., the maximum link ratio between the two best contig pairs; default = 0.3), which were tested for all combinations of *l*=5-10, *a*=0.1-0.9. Optimization focused on increasing N50 but also balancing the number of misjoined scaffolds observed in the third stage of scaffolding with Hi-C (*see below*). The selected parameters (i.e., *l* = 8; *a* = 0.2) had a maximum of two or three visible misjoins. This stage introduced some short contigs (< 200 bp) assembled within scaffolds without meaningful gap sizes, and these were removed using *sed*, with all remaining gaps resized to strings of 100 Ns. The small number of scaffolds that were smaller than 1 kbp (i.e., fewer than 100 scaffolds) were removed using *fastx* of Bioawk (Li, 2017).

In the third scaffolding stage, Hi-C data was re-aligned to the genome following the scaffolding stages above using Juicer v.1.5.6 (Durand, Shamim, et al., 2016) with flags and arguments *-s* Sau3AI (i.e., the restriction enzyme applied) and *-y* to include the restriction site file, as well as *-S* early to indicate early exit of the program. The resultant output (i.e., merged_nodups.txt) was used with 3d-dna v.180922 (Dudchenko et al., 2017) using parameters *-i* 50000 (i.e., the minimum size of contigs to scaffold) and *-r* 0 (the number of iterative rounds for misjoin correction). The assembly was visualized after scaffolding using Juicebox v1.8.8 (Durand, Robinson, et al., 2016) to identify and split mis-assemblies, and to identify, order, and orient linkage groups (LGs). LG-like groups were oriented such that the greatest density of inter-chromosomal contacts was at the 5’ end of each LG. Using *juicebox_assembly_converter.py* (Phase Genomics), NCBI AGP files were generated. The newly generated LGs were compared to the northern pike v.1.0 assembly (Rondeau et al., 2014) using LastZ (Harris, 2007) visualized in Geneious (Kearse et al., 2012) using default parameters, and the AGP file was manually edited to keep naming consistent to the v.1.0 assembly. The genome was then submitted to NCBI (see *Data Availability*).

Genome annotation was conducted using the NCBI Eukaryotic Annotation pipeline v.8.2 under annotation release 103. Genome completeness was evaluated using BUSCO v.4.0.2 with both Actinopterygii and Vertebrata odb10 datasets (Seppey et al., 2019). Analysis of repeat content used the same methods and custom repeat library as previously described (Rondeau et al., 2014).

### Population genomics: sampling, sequencing, and analysis

Northern pike tissue samples (n = 47) were obtained from across Canada and the northern United States as provided by collaborators and hatcheries (see Table 1; Figure 1). Fish from the upper St. Lawrence River were collected and processed following a protocol approved by the SUNY ESF Institutional Animal Care and Use Committee. Alaskan Minto Flats samples used for sex marker validation (*see below*) were provided as fin clips in 95% ethanol by the Alaska Department of Fish and Game following approved state and departmental regulations and protocols. All other tissues were either archival, opportunistic sampling of fishery harvest or government purposes (e.g., invasive species control), and therefore did not require ethical review by the University of Victoria in accordance with the Canadian Council on Animal Care Guidelines on the care and use of fish in research, teaching and testing (s4.1.2.2; CCAC, 2005).

**Figure 1.**
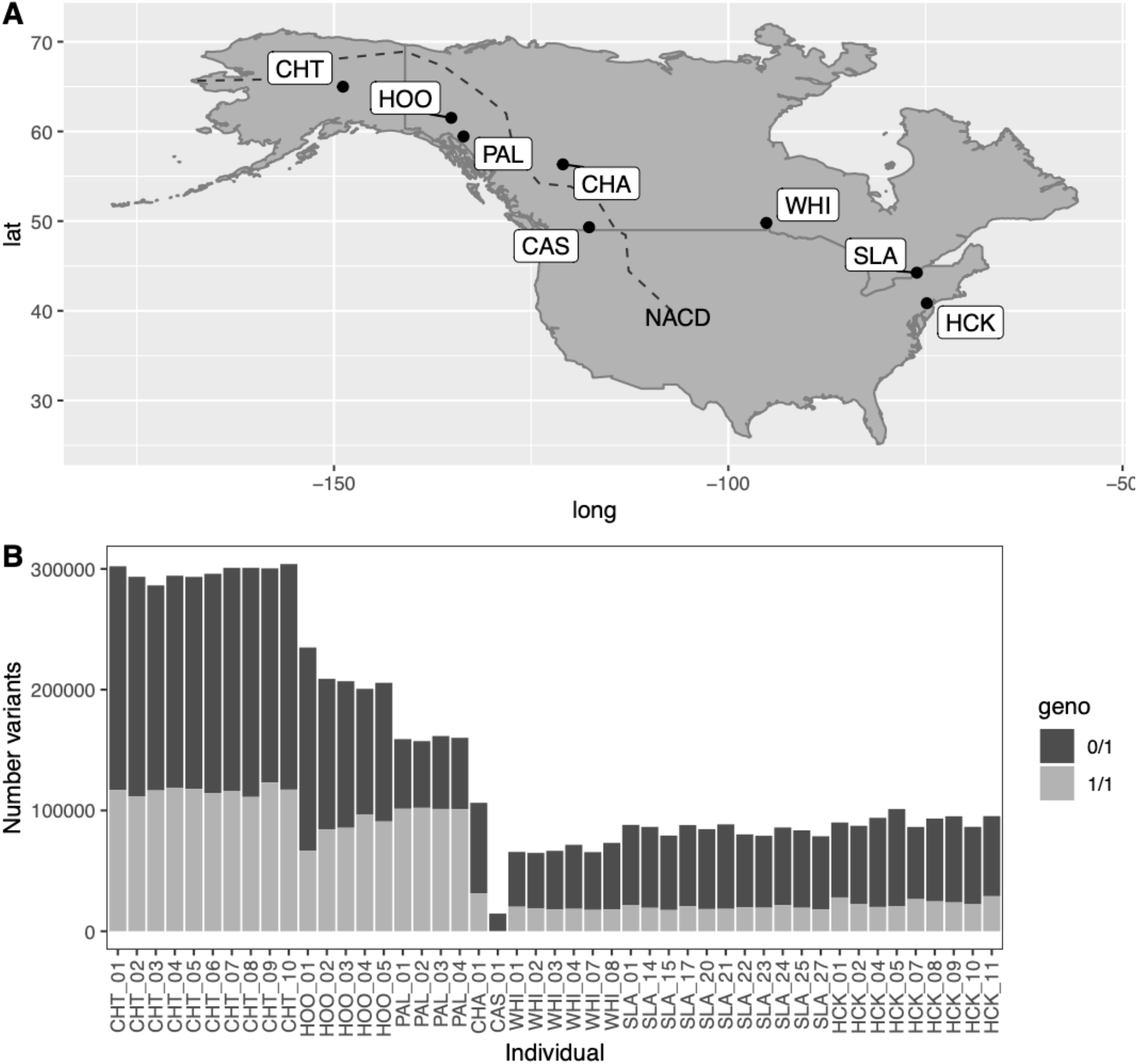
(A) Sampling locations for northern pike across North America, including Chatanika River, Alaska (CHT), Yukon River at Hootalinqua, Yukon Territory (HOO), Palmer Lake, British Columbia (B.C.; PAL), Charlie Lake, B.C. (CHA), Columbia River at Castlegar, B.C. (CAS), Whiteshell Hatchery, Manitoba (WHI), St. Lawrence River at New York (SLA), Hackettstown Hatchery, New Jersey (HCK). (B) Per individual total counts of homozygous alternate (light shading) or heterozygous (dark shading) variants.

**Table 1.** Sampling location details and sample sizes classified using phenotypic sex information. Sample sites with unrecorded phenotypic sex are indicated as unknown (*unkn*.). The most recent genome assembly (v.4.0) was constructed using the Castlegar female. Samples were obtained throughout North America including from Alaska (AK), the Yukon Territories (YT), British Columbia (BC), Manitoba (MB), New York (NY), and New Jersey (NJ). Populations obtained from hatcheries are indicated with an asterisk. The short names for each sample site are provided.

| Country | Prov/<br>State | Population | Short<br>name | Latitude | Longitude | Mal.<br>(n) | Fem.<br>(n) | Tot.<br>(n) |
| --- | --- | --- | --- | --- | --- | --- | --- | --- |
| USA | AK | Chatanika R. | CHT | 64.98396 | -148.86032 | 5 | 5 | 10 |
| Canada | YT | Hootalinqua | HOO | 61.51156 | -135.13209 | *unkn.* | *unkn.* | 5 |
| Canada | BC | Palmer Lk. | PAL | 59.43708 | -133.57592 | *unkn.* | *unkn.* | 4 |
| Canada | BC | Charlie Lk. | CHA | 56.32853 | -120.97835 | 1 | - | 1 |
| Canada | BC | Castlegar | CAS | 49.31538 | -117.65344 | - | 1 | 1 |
| Canada | MB | Whiteshell | WHI* | 49.80051 | -95.17243 | 3 | 3 | 6 |
| USA | NY | St. Lawrence | SLA | 44.24787 | -76.09785 | 6 | 5 | 11 |
| USA | NJ | Hackettstown | HCK* | 40.84155 | -74.83359 | 6 | 3 | 9 |
| **Total** |  |  |  |  |  | 21 | 17 | 47 |

DNA was extracted from a variety of tissues using DNEasy Blood and Tissue Kit (QIAGEN) following the manufacturer’s protocols, then quantified by Nanodrop ND-1000 (ThermoFisher) and Qubit v2.0 (Life Technologies). Samples were sent to McGill University and Genome Quebec Innovation Centre for library preparation and sequencing. Most of the samples (i.e., 35 of 47) underwent PCR-free whole genome shotgun sequencing. The ten samples from the Chatanika River had insufficient quantities for PCR-free libraries and therefore were sequenced via PCR shotgun sequencing. All 45 of these libraries were sequenced on an Illumina HiSeqX Ten in paired-end 150 bp reads with between five and seven samples sequenced per lane. The two remaining samples (i.e., Castlegar and Charlie Lake, BC) were used for reference genome assembly, with the Castlegar individual being the focus for the final v.4.0 assembly. Due to the focus on detecting sex determination loci, lanes were designated as sex specific as much as possible to reduce the impact of potential index switching between sexes.

Read processing and variant calling was based on GATK’s best practices (GATK v.3.8-0-ge9d806836; DePristo et al., 2011; McKenna et al., 2010; Poplin et al., 2018; Van der Auwera et al., 2013). In brief, paired-end reads were aligned to the latest northern pike genome (*described above*; GCF_004634155.1) using bwa-mem v.0.7.13-r1126 (Li, 2013). Alignment files were converted to bam format, sorted, and indexed by position using SAMtools v.1.3. Sequencing platform and multiplexing layout metadata was used to mark duplicates using Picard v.2.17.11 to flag for downstream genotyping. The two samples used for reference genome assemblies (i.e., Castlegar and Charlie Lake, BC) had 5-7x deeper coverage than the rest of the samples, and so they were down-sampled (i.e., reads were randomly subsampled to the targeted coverage of 25X for Charlie Lake and 20X for Castlegar) using SAMtools view with the *-s* flag. Nucleotides in all bam files were re-calibrated according to GATK’s recommendations for non-model organisms.

Variants were called from re-calibrated bam files independently per sample using the *HaplotypeCaller* of GATK in GVCF mode, then combined as a cohort using the *GenotypeGVCF* function of GATK to produce a VCF file containing 1,910,789 SNPs and insertion/deletions (indels) for all 47 samples. The 1,363,731 SNPs within this VCF were then extracted and filtered (see Table 2). A hard filter was applied using GATK to remove variants according to the following cutoffs or parameters: quality by depth = 2; fisher strand bias = 60; root mean square mapping quality = 30, mapping quality rank sum test = -12.5; and read position rank sum test = -8.0. Additional quality control filters were applied using VCFtools v.0.1.15 (Danecek et al., 2011) including removing sites where more than 10 individuals had missing data (--*max-missing-count* 10) or those with no minor alleles observed (*--mac* 1). Sites were further required to have at least one homozygous call, either reference or alternate. The resultant VCF file with 1,127,943 variants was used for the downstream analysis, although further filtration was conducted for specific analyses as discussed in individual sections below.

**Table 2.** Variant filtering steps and parameters applied after alignments of reads against the northern pike reference genome (v.4.0).

| Filter Step | Parameter<br>value | Program | No. variants<br>retained |
| --- | --- | --- | --- |
| Raw genotypes<br>(includes indels) | - | - | 1,910,789 |
| SNP variants only | - | GATK | 1,363,731 |
| GATK hard filter | - | GATK | 1,189,068 |
| Min. quality | 20 | VCFtools | 1,186,793 |
| Min. mean depth | 10 | VCFtools | 1,153,668 |
| Max. mean depth | 60 | VCFtools | 1,152,122 |
| Max. missing count | 10 | VCFtools | 1,129,884 |
| Minor allele count | 1 | VCFtools | 1,129,701 |
| Per locus heterozygous<br>genotypes | < 100% | R/ VCFtools | **1,127,943** |

The numbers of variant calls per individual were summarized and visualized in R (R Core Team, 2023) using vcfR v.1.14.0 (Knaus & Grünwald, 2017) and dplyr v.1.1.4 (Wickham et al., 2023). Genotype counts per site were obtained using the *VariantsToTable* function of GATK for heterozygotes, homozygous reference, homozygous variant, and no call, as well as the total number of variants and samples called per site. To obtain per site genotype frequencies, each category was divided by the number of samples called. Observed heterozygosity (*H_OBS_*) was calculated by reading in the VCF into R, converting it to genind format using vcfR then to genlight format to calculate *H_OBS_* with the function *gl.report.heterozygosity* using dartR v.2.9.7 (Gruber et al., 2018). Private alleles per population were calculated using the *private_alleles* function of poppr v.2.9.4 (Kamvar et al., 2014). Tajima’s D in bin sizes of 10,000 were calculated using bcftools (Danecek et al., 2021).

A maximum likelihood tree based on genome-wide SNPs was generated using SNPhylo v.20140701 (Lee et al., 2014) using default parameters and with 1,000 bootstraps. The resulting tree was visualized using FigTree version 1.4.3 (Rambaut, 2016), and rooted by midpoint. A discriminant analysis of principal components (DAPC) was performed with bi-allelic genome-wide SNPs using adegenet v.2.1.1 (Jombart, 2008; Jombart & Ahmed, 2011) in R. The *find.clusters* function of adegenet was used to determine the appropriate number of clusters based on the lowest Bayesian Information Criterion value when all principal components were kept, and DAPC was conducted on these groups, retaining 24 principal components and three discriminant functions. The *snpzip* function with the Ward clustering method was used to list SNPs with the greatest contribution to each of the three discriminant axes identified.

### Characterizing diversity across chromosomes

The genome assembly was unwrapped using custom python script *fasta_unwrap.py* (E. Normandeau, Scripts; see *Data Availability*) and subset to only contain chromosomes. Chromosome lengths were calculated using *fasta_lengths.py* (E. Normandeau, Scripts). The genome was then indexed using samtools, and a bed file containing 1 Mbp windows was prepared using the *makewindows* function of bedtools. The number of variants per window for each chromosome was then calculated using the *coverage* function of bedtools. The distribution of variants per window for all windows was then used to determine the minimum number that would be considered an outlier (i.e., third quartile + 1.5x the interquartile range) using the *boxplot.stats* function of R, and the number of variants per window and the mean number of variants per kbp per chromosome were plotted using custom scripts in R (see *Data Availability*).

### Empirical sex-linked variants and genotypes in resequencing data

The identification of k-mers associated with sex was performed on the resequenced populations that had at least three females and three males (i.e., Chatanika River (CHT), Manitoba (WHI), New Jersey (HCK), and New York (SLA); see Table 1). Raw reads were concatenated and then all possible 31mers were extracted to form a master list of k-mers using Jellyfish v.2.2.6 (Marçais & Kingsford, 2011) running on Compute Canada. The master list was then used to query the reads of each individual sample using Jellyfish to count all present 31mers. A sex-specific analysis was then conducted whereby females and males were compared within each population. Sex-specific k-mers were considered as those sequences for which all of the opposite sex in the population had two or fewer copies (to allow for sequencing errors), and the target sex individuals all had at least one k-mer and the sum of all males had more than seven instances of the k-mer. The resulting sex-specific k-mer sets for each population were mapped back to the reference genome using bwa-aln v.0.7.13-r1226 (Li, 2013). Following the alignment, the number of sex-specific k-mers per population were counted in 10 kb windows across the genome using bedtools v2.26.0 (Quinlan & Hall, 2010) and plotted in R v.3.5.3 using ggplot2 (Wickham, 2016).

Second, a genome-wide association analysis was performed using sex as the phenotype. The individuals with sex phenotype data were extracted from the VCF. This extraction also included a filter to keep variants with a minor allele count of at least two (i.e., *--mac* 2), resulting in the analysis of 17 females and 21 males. The analysis was performed using plink v.1.9b_5.2-x86_64 (Purcell et al., 2007) using the *fisher-midp* option to run Fisher’s exact tests and output p-values. The -log(p-value) of each SNP was visualized in a Manhattan plot using qqman v.0.1.4 (Turner, 2014). Significant associations were considered when Bonferonni corrected p ≤ 0.05.

Third, a DAPC was conducted using sex as the differentiating variable using all resequenced individuals with sex phenotypes within populations and defined groups from PCA and DAPC (*see above*) in adegenet. Population and group-specific SNPs were extracted from the VCF file. Each group was filtered independently using a custom script in R to remove variants that were not expected to be related to the sex determination system (i.e., homozygous alternate variants). These variants were removed because the reference genome is female and the sex determination system expected to be XY, and therefore the homozygous alternate variants were not expected to be part of this system. The remaining sites were those with homozygous reference and heterozygous genotypes. Subsequently, the DAPC analysis was performed on each subset. Output loadings tables were inspected for genotypes where all males or all females were heterozygous. From this information, lists of genomic locations with putatively sex-specific signatures were generated for each population. Histograms of sex-specific SNP occurrences were then generated along LGs using ggplot2.

### Sex markers and histology

PCR-based inspection for genetic sex markers was conducted for individuals used in resequencing (*see above*), as well as additional samples obtained from Castlegar, BC (n = 2 females and 5 males) and from the Minto Flat region of Alaska, USA (n = 6 females, 14 males, 1 undetermined). PCR used primers designed to amplify regions of *amhby* (i.e., SeqAMH1-4 and ConserveAMH1-1; Pan et al., 2021), as well as newly developed primers from this study (i.e., set_24.5; Table S1). The set_24.5 primers amplify a 500 bp amplicon in both sexes, and an additional 250-300 bp amplicon specifically in genetic males using a nested primer pair based on male-specific SNPs. PCR was conducted using 2 mM MgCl_2_, 0.2 mM dNTPs, 1X reaction buffer, 0.25 units of Taq polymerase (Promega), 0.5 µM of each primer, 50-100 ng of DNA, a remaining volume of molecular grade water up to 10 µl total reaction volume. The nested primer pair used 0.2 µM forward primer instead of 0.5 µM and included 0.3 µM of the probe. All PCR used the following thermal regime: 95°C for 5 min., 35 cycles of 95°C for 0.5 min., 0.5 min. anneal, 72°C extension for 1 min. (or 0.5 min. for set_24.5), and a final 10 min. extension at 72°C. PCR products were visualized on 1% agarose gels and scored as female, male, or undetermined.

Histological determination of phenotypic sex was conducted using gonadal tissue from the Minto Flats and the St. Lawrence River populations. In brief, Minto Flats samples were saturated with a 1:1 solution of 100% ethanol:LR White resin (hard grade) for 24 hours, then with pure LR White for 24 hours (resin replaced fresh after 6 hours). Each sample was then placed in separate gelatin capsules with fresh catalyzed LR white and polymerized at 60°C for 24 hours. Samples were cut to 1-micron sections, stained with Richardson’s LM stain, and examined microscopically. St. Lawrence River samples were preserved in Davidson’s solution for 48 hours, transferred to 70% ethanol prior to histological processing as described above. Following processing, tissues were embedded in paraffin wax and 5-micron sections were cut and stained with hematoxylin and eosin (H&E).

## Results

### Genome assembly and annotation

A northern pike female was sampled from the Canadian portion of the Columbia River at Castlegar, British Columbia (BC) and DNA was extracted from liver tissue to preserve high molecular weight genomic DNA. The DNA from this individual was sequenced to approximately 80X depth using 8 SMRT cells on a Pacific Biosciences Sequel instrument. The initial contig-level assembly generated by Canu (see *Methods*) yielded a total length of 939.0 Mbp in 1,258 contigs (contig N50: 3.9 Mbp). Polishing and scaffolding occurred in several stages as described in brief here (see detailed description in *Methods*). First, Hi-C data was applied to the assembly resulting in 941 scaffolds (scaffold N50: 18.8 Mbp). Second, 10X Chromium data was applied to the assembly, which split problematic contigs and allowed for additional scaffolding. The error correction step increased the number of contigs to 1,395 and reduced contig N50 (contig N50: 3.4 Mbp), but the additional scaffolding increased the scaffold N50 (scaffold N50: 23.3 Mbp). Third, the Hi-C data was further applied and manually reviewed to yield a total of 811 scaffolds (final scaffold N50: 37.6 Mbp; Figure S1). This final northern pike assembly (v.4.0) is 941 Mbp in length, with a maximum scaffold length of 52.6 Mbp, and includes 25 scaffolds of the expected chromosome lengths. In addition, the assembly contains 785 unplaced scaffolds with a total length of 23 Mbp.

The 25 chromosome-length scaffolds were then assigned to the 25 linkage groups of northern pike (Rondeau et al., 2014). Linkage groups were oriented by density of inter-chromosomal contacts, where the greatest density of repeat elements was oriented to the 3’ end of the chromosomes (Figure S2). This fits with the original orientation by Rondeau and co-workers (2014), but should be noted that the centromeres for the linkage map and assembly are therefore positioned at the 3’ ends of all LGs or chromosomes. A full inventory of observed repeat elements and their percentages is shown in Table S2. The mitochondrial genome was identified by BLAST alignment of the previous assembly (NC_025992.1; Rondeau et al., 2014) against the present assembly. Once identified, the mitochondrial genome was manually circularized. The v.4.0 assembly has been submitted to NCBI under assembly accession GCA_004634155.1 (see *Data Availability*).

Annotation of the assembly was performed by the NCBI eukaryotic genome annotation pipeline using RNA-sequencing data (Pan et al., 2019; Rondeau et al., 2014) and EST data (Leong et al., 2010). This identified 24,843 protein-coding genes. A BUSCO analysis found that 95.4% and 97.1% of the Actinopterygii (odb10) and Vertebrate (odb10) genes were complete, respectively. Relative to the former database with 3640 genes, 94.1% (i.e., 3426) were identified as single copies, and only 1.3%, 1.3%, and 3.3% were duplicated, fragmented, or missing, respectively.

### Genetic variation and population genomics

Northern pike were obtained for whole-genome resequencing from across North America (Figure 1; Table 1). Samples included those from Chatanika River (CHT) in Interior Alaska, USA (n = 10), Yukon River at Hootalinqua (HOO) in the Whitehorse Region of the Yukon Territories, Canada (n = 5), Palmer Lake (PAL) in northwestern British Columbia (BC; n = 4), Charlie Lake (CHA) in northeastern BC (n = 1), Columbia River at Castlegar (CAS) in southeastern BC (n = 1), Whiteshell Hatchery (WHI) in Manitoba (n = 6), the St. Lawrence River (SLA) off New York State (n = 11), and at Hackettstown Hatchery (HCK) of New Jersey, USA (n = 9). In total, this included 17 phenotypic females, 21 males, and nine individuals with undetermined sex (n = 47 total; Table 1). The individual sampled at Charlie Lake and the individual sampled at Castlegar were used for whole-genome sequencing, and the remainder of the samples were used for whole-genome resequencing (see *Data Availability*). The Charlie Lake assembly was not used in this study, but the sample’s data was used in the rest of the analyses as a representative for this collection site.

In total, 1,363,731 raw SNP variants were called in the whole-genome resequencing data, and after all quality filtering, 1,127,943 SNPs were retained (i.e., 82.7% retention; see Table 2). On average, the mean (± s.d.) depth per individual per variant site was 20 ± 3.3x (range = 15x to 30x). There was a large difference in genetic variation observed across sampling sites (Figure 1B; Table 3). Interior Alaska samples (CHT) had the highest rate of heterozygous genotypes (mean per individual = 180,943 SNPs), followed by the Yukon River (HOO; 126,720 SNPs). All other populations had fewer heterozygous SNPs, with 58,129 heterozygous SNPs in northwestern BC (PAL), 49,401 in Manitoba (WHI), 64,394 in New York (SLA), and 67,944 in New Jersey (HCK). Additionally, Alaska, Yukon, and northwestern BC had similarly high numbers of homozygous alternate genotypes (mean per individual: 116,276, 84,775, and 101,465, respectively), and the eastern BC and eastern Canada/US samples had similarly lower numbers of homozygous alternate genotypes ranging from 18,525 to 31,385 on average per individual (Additional File S1). Considering the ungapped genome length of 940.8 Mbp, and including both heterozygous and homozygous alternate variants, the CHT and SLA populations have an average per individual difference from the reference genome of 0.0316% and 0.00891%, respectively. In terms of the average SNP per kbp per individual, CHT and SLA have a SNP for every 3,165 bp and 11,222 bp, respectively. The average observed heterozygosity (*H*_OBS_) per population followed a similar trend as heterozygous SNP counts, with *H*_OBS_ of CHT and HOO estimated at 0.161 and 0.113, respectively, whereas PAL, WHI, SLA, and HCK ranged from 0.044-0.060. More pronounced negative Tajima’s D values were observed in WHI and SLA, with values of -1.07 and -1.30, respectively, than HCK (-0.33) or the higher diversity populations CHT and HOO (-0.22 and -0.331, respectively; Table 3). Results for the populations with only a single individual should be taken with caution given the low sample size, different technology applied to sequencing, and for the Castlegar sample (CAS), being the reference genome used in the study.

**Table 3.** Per population summary statistics following genotyping, including the average number of heterozygous or alternate homozygous variants per individual, the per population observed heterozygosity, Tajima’s D, and the number of private alleles. Sample size was rounded to nearest whole number. Acronyms: CHT = Chatanika R.; HOO = Hootalinqua; PAL = Palmer Lake; CHA = Charlie Lake; CAS = Castlegar; WHI = Whiteshell; SLA = St. Lawrence River; HCK = Hackettstown.

| Pop | (n) | Average no.<br>variants<br>per indiv.<br>(ref/alt) | Average no.<br>variants<br>per indiv.<br>(alt/alt) | Obs.<br>heterozyg.<br>($H_{OBS}$) | Tajima's<br>D | Private<br>alleles |
| --- | --- | --- | --- | --- | --- | --- |
| CHT | 10 | 180,943.2 | 116,275.7 | 0.1609 | -0.221 | 204,410 |
| HOO | 5 | 126,719.8 | 84,774.6 | 0.1127 | -0.331 | 67,803 |
| PAL | 4 | 58,128.5 | 101,464.5 | 0.0516 | -0.072 | 21,290 |
| CHA | 1 | 74,996.0 | 31,385.0 | 0.0668 | - | 14,736 |
| CAS | 1 | 14,552.0 | 96.0 | 0.0129 | - | 17,974 |
| WHI | 6 | 49,401.8 | 18,524.8 | 0.0438 | -1.068 | 101,149 |
| SLA | 11 | 64,393.5 | 19,442.5 | 0.0571 | -1.300 | 214,742 |
| HCK | 9 | 67,943.5 | 24,188.0 | 0.0604 | -0.328 | 107,085 |

Genetic similarity between individuals and populations based on clustering in a genetic dendrogram followed a similar trend to that observed in overall variation (i.e., considering the sum of both heterozygous and homozygous alternate genotypes), where the Alaska (CHT), Yukon (HOO), and northwestern BC (PAL) populations clustered separately from all other samples (Figure 2). Most of the populations had very strong support (100%) for grouping all individuals in the population together, and only the Yukon (HOO) population did not show a highly supported single cluster. Within each separate section of the dendrogram for high or low diversity samples, the closest populations geographically were also the most proximal in the dendrogram.

**Figure 2.**
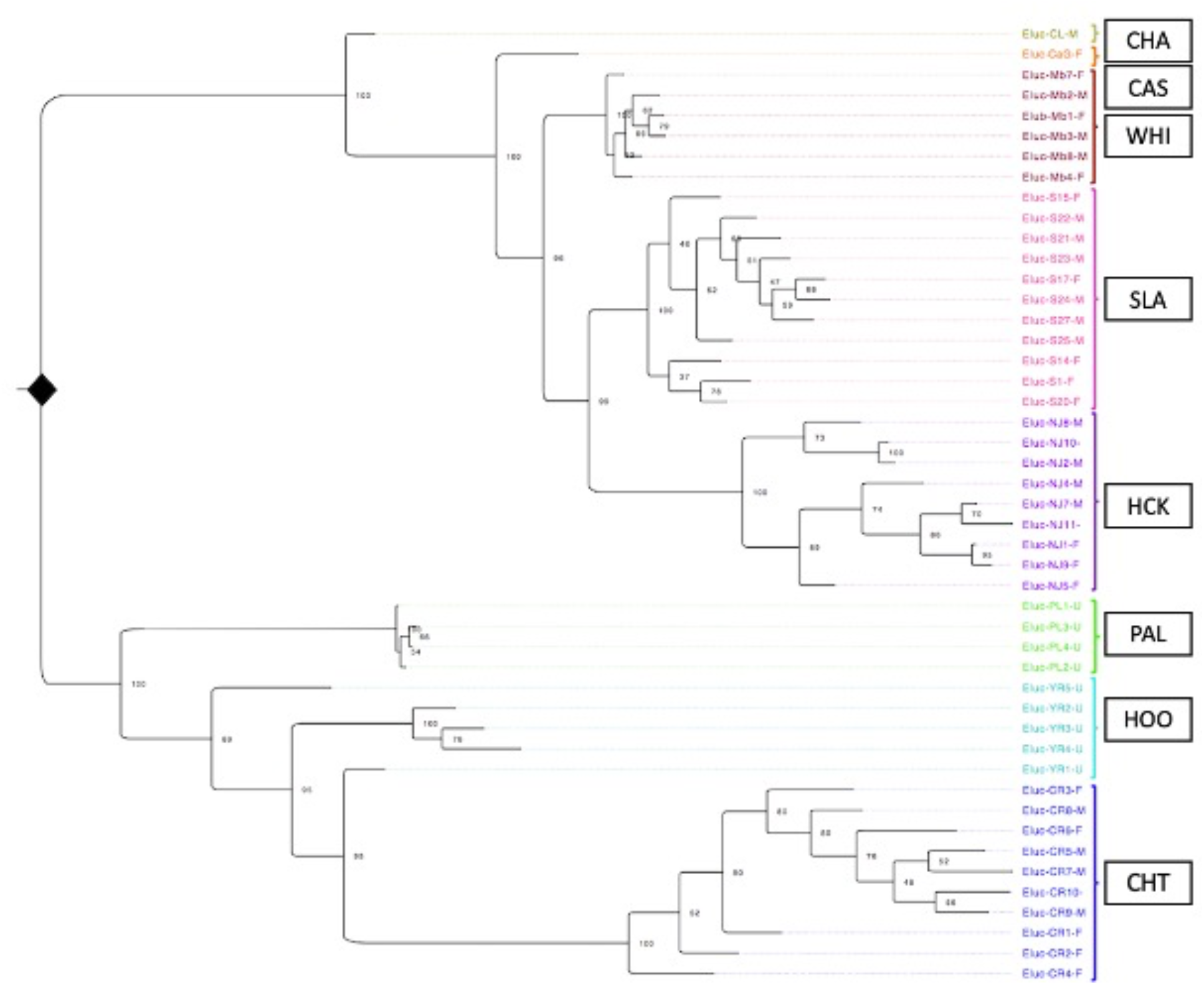
Genetic dendrogram clustering of individual northern pike using all filtered SNP variants. Bootstrap support is indicated at branch nodes as evaluated by 1,000 bootstraps. The largest separation in the data separates samples across the North American Continental Divide (NACD), with populations from Alaska (CHT), Yukon (HOO), and northwestern British Columbia (PAL) separating from all other populations. See Table 1 for all acronyms.

An analysis of private alleles and fixed differences further indicates the genetic separation and isolation of the different populations. Of the 1.1 M variants characterized, 0.75 M were only present in a single population (i.e., private alleles), and although many are of low frequency, each population with at least four individuals had between 21,290-214,742 private alleles present. As an example, CHT has 81,263 private alleles with at least four instances of the allele being observed, whereas HOO has 5,395, HCK has 26,173 and SLA has 5,989 private alleles with at least four instances of the allele. Furthermore, when considering fixed homozygous alternate genotypes, a large proportion of these were unique to each population. For example, there were 34,561 biallelic variants that were observed as only homozygous alternate genotypes in Alaska (CHT), and of these, 34,002 were fixed reference in St. Lawrence River (SLA) and 33,285 were fixed reference in the Yukon (HOO). As another example, there were 13,777 biallelic variants observed as only homozygous alternate genotypes in SLA, and of these, 10,975 and 9,993 were observed as fixed homozygous reference in the Hackettstown (HCK) or Alaska (CHT) populations, respectively. Therefore, many variants identified here were specific to one of the populations, and many variants were fixed in specific populations, even in geographically proximal populations.

The distribution of variants was not equal across the northern pike genome (Figure 3A). When considering the sum of variants within 1 Mbp windows across chromosomes (n = 930 windows total), 59 windows were considered outliers in terms of elevated polymorphism (see *Methods*) with more than 2,341 variants per window in comparison with the genome-wide average of 906 variants per window (median: 1,202; Additional File S2). Several of the chromosomes have large sections of elevated variation where multiple consecutive windows are outliers, specifically chromosomes 5 (3’ end, 24-29 Mbp), 9 (5’ end, 4-10 Mbp), 11 (3’ end, 35-44 Mbp, excluding 41-42 Mbp), and 24 (throughout, 0-5 Mbp excluding 3-4 Mbp, and 18-23 Mbp; Figure 3A). Furthermore, when considering a per-chromosome average number of variants per kbp, chromosome 24 has a clear elevation overall in terms of polymorphism level relative to all other chromosomes (Figure 3B). Notably, in populations that still use *amhby* as the sex determining gene, this gene is in chromosome 24. Some preliminary exploration of elevated polymorphic regions suggests presence of genes related to immunity or having multiple copy numbers, for example chromosome 09 contains zinc-finger genes, immune-associated nucleotide-binding protein genes (GIMAPs), as well as multiple genes of the major histocompatibility complex (MHC) class II type, but it is beyond the scope of this paper to describe all of the regions of the northern pike genome with elevated SNP density.

**Figure 3.**
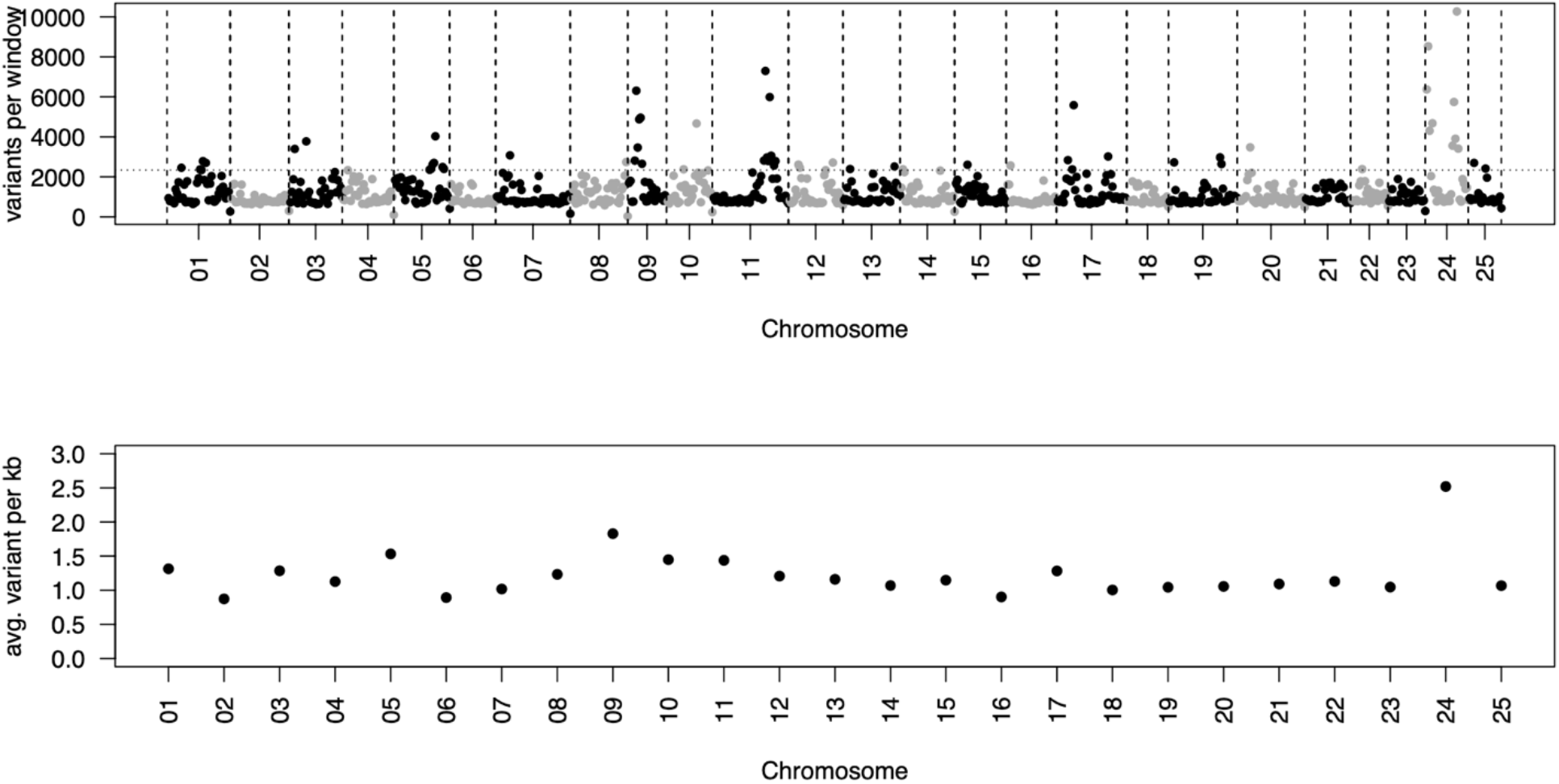
(A) Genome-wide SNP density by chromosome showing all SNP variants retained in the dataset, with variants summed across 1 Mbp windows (n = 930 windows total). Windows above the outlier threshold are indicated by the horizontal hatched line (*see Methods*). (B) The per chromosome average number of variants per kbp are shown, indicating chromosome 24 as a clear outlier overall.

### Sex determination

The dataset was restricted to only include individuals with phenotypic sex recorded, and following filters, there were 672,565 SNPs retained across 17 females and 21 males. All individuals were initially analyzed together (i.e., on a continental scale) using three different approaches to search for a sex-specific locus in the data. Both a genome-wide association study (GWAS) and a DAPC with sex as the separating variable were performed but found no significant or suggestive associations (*data not shown*). Additionally, a k-mer analysis of raw sequence reads and therefore operating independently of a reference genome also failed to identify sex-specific polymorphism at this scale. The reference-independent approach was valuable given that the reference genome used to score SNPs was developed from a female individual, and therefore the male-specific sex determining locus would be expected to be missing (Pan et al., 2021).

After the continental-scale analyses failed to find sex-associated loci, population-specific analyses were conducted. With this focused analysis, a sex-specific signal was detected in the Alaskan sample (CHT; n = 5 individuals per sex), where male-specific heterozygosity was observed on LG24 through both the DAPC and k-mer analyses (Figure 4), as previously observed (Pan et al., 2019). In the CHT samples, a 500 kb region between 650-1150 kb of LG24 (total chromosome length: 29.5 Mbp) was detected, containing 3,552 male-specific SNPs. In addition, 1,137 male-specific and 39 female-specific SNPs from other sections of the genome were also identified by DAPC analysis (Additional File S3). Although none of the Yukon River (HOO) samples had associated phenotypic sex information, one of the five individuals also held the male-specific SNPs on LG24 (and elsewhere) identified from the Alaska population on LG24. However, all other populations assessed in North America did not hold these sex-specific variants (Figure 4).

**Figure 4.**
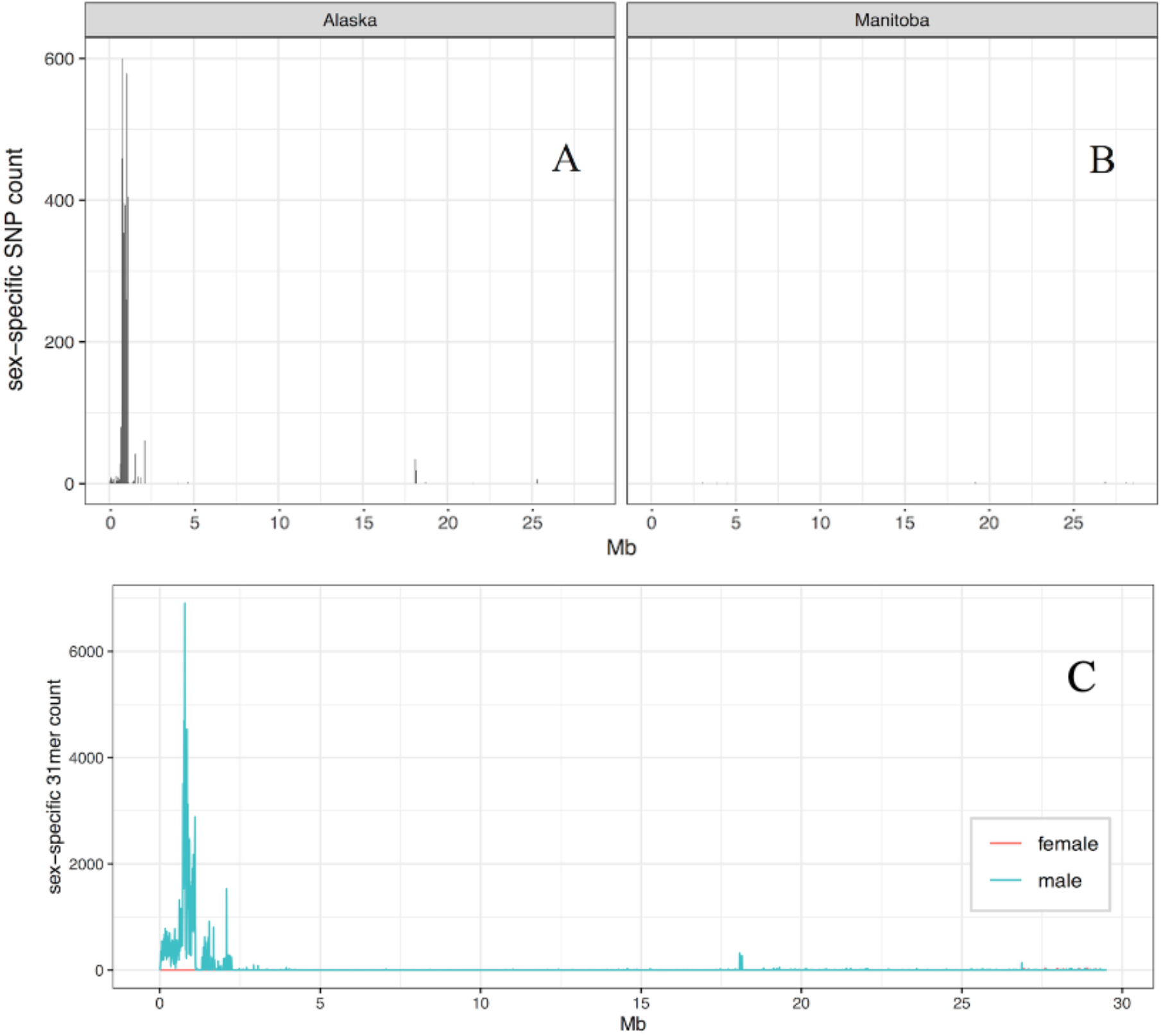
Sex-specific SNP counts for LG24 shows a strong signal in the Alaska population (panel A; CHT), and a lack of signal for populations east of the NACD (panel B; WHI used as example). The sex-specific signal is also observable in the Alaskan LG24 when viewing k-mer content rather than variant content. The k-mers were identified independent to the reference genome, but were aligned to the genome after identification.

To further characterize the genetic sex of the sampled individuals across North America, since the reference genome is generated from a female and therefore is not expected to have the male-specific sex-determining locus *amhby*, the presence of *amhby* was investigated using *amhby*-specific primers in a PCR analysis (Pan et al., 2021). This confirmed that *amhby* was present in the phenotypic males from Alaska (CHA) and the putative male from the Yukon (HOO) as observed in the sequence data. None of the other samples in any collection used for WG-resequencing in North America amplified *amhby*. The PCR results therefore agree with the lack of a sex-specific signal in the other populations by GWAS, DAPC, or k-mer analyses (*see above*).

Additional sampling was undertaken to further characterize the geographic regions in which the transition from *amhby*(+)-males to *amhby*(-)-males occurs. Two additional females and three males were obtained from the Columbia River at Castlegar, BC (CAS), and six females and 14 males were obtained from Minto Flats, Alaska. These samples were tested for *amhby* by PCR markers (Pan et al., 2021), as above. Interestingly, all three of the Castlegar males were positive for *amhby* (females were negative); this contradicts the expectation of the absence of *amhby* in males in this region. Also unexpectedly, the Minto Flats males, expected to all be *amhby*(+), did not show a uniform positive detection with only six of the 14 phenotypic males with a positive *amhby* detection (the females were all negative). All three primer pairs, including *amhby_conserve* (partial exon 2) and *seqAMH_1* (partial exon 7) from Pan et al. (2021), and primer pair set_24.5 (LG24: 996,878-997,339 bp) found concordant results for these samples. Sex phenotypes were further evaluated through gonadal histology, which confired testes development in both *amhby*(+) and *amhby*(-) males (Figure 5). Therefore, there are both *amhby*(+) and *amhby*(-) males in Alaska, and *amhby*(+) males further southeast than expected in North America.

**Figure 5.**
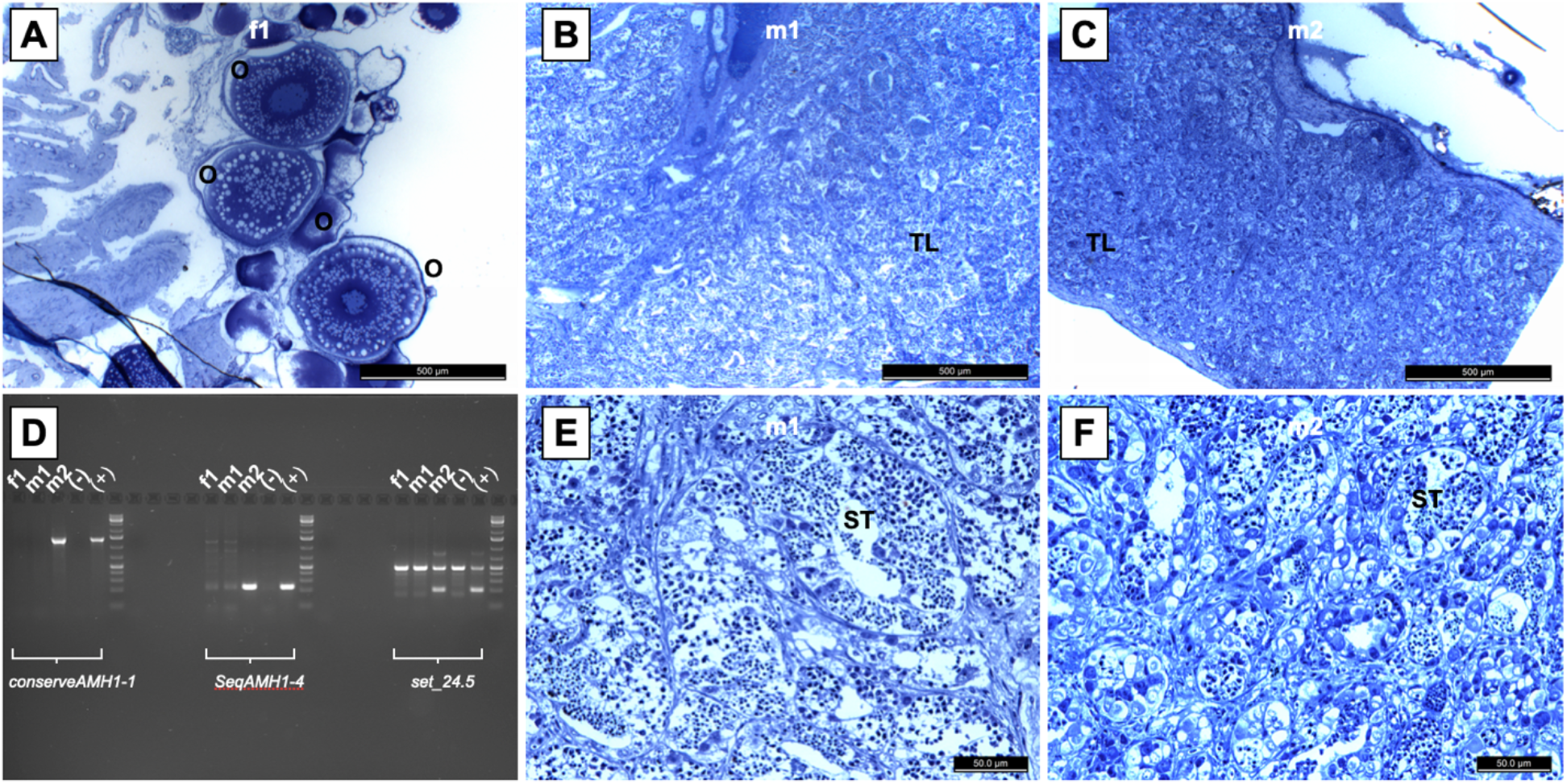
Gonadal histology and *amhby* PCR testing for sex genotypes and phenotypes in additional samples from Minto Flats, Alaska. The phenotypic female (f1) is shown in panel (A) with ooctyes (O) indicated at various stages of vitellogenesis. The two phenotypic males m1 and m2 are shown in panels (B) and (E), and panels (C) and (F), respectively, with testis lobules (TL) and seminiferous tubules (ST) indicates with gametes at various stages of spermatogenesis. In panel (D), PCR results of amplicons developed with the three primer pairs applied (see *Methods*) for the female (f1), an amhby(-) male (m1), and an amhby(+) male (m2), as well as a negative and positive control sample for each primer pair, selected from samples from Alaska (CHT) in the resequencing data. Primer sets *conserveAMH1-1* and *SeqAMH1-4* by Pan et al. (2021) indicate *amhby* by a single band, and *set_24.5* indicates presence of *amhby* by a second band.

## Discussion

### Genetic diversity across North America

In all the populations assessed across North America from Alaska through New Jersey, the northern pike from the west in Interior Alaska and the Yukon River have the highest genomic diversity. A striking decrease in overall diversity coincides with the location of the North American Continental Divide (NACD; i.e., the separating point between eastward or westward-flowing watersheds). West of the NACD, a stepwise decrease in diversity occurs from Interior Alaska to the Yukon, then to northwestern British Columbia, as measured by observed heterozygosity and the total number of heterozygous variants. East of the NACD, the populations have uniformly low diversity. The eastern region samples at the Manitoba hatchery (Whiteshell) and the St. Lawrence River had more pronounced negative Tajima’s D values, suggesting that these populations were reduced in number and then more recently expanded, resulting in an excess of low frequency polymorphism (Hedrick, 2005; Tajima, 1989). This was not universally observed in the eastern samples (e.g., Hackettstown had a similar Tajima’s D value as western sites). The NACD may not be causal to the diversity decrease, although it is an important land formation that influences contemporary water catchments and flow, but it does serve as a marker beyond which all populations surveyed to date have been found to have universally low diversity. The west to east diversity decrease is in agreement with theory that the western populations are the oldest in North America, and that while these populations expanded eastward from the Beringia refugium, for example while recolonizing post-glacial recession, they were impacted by population bottlenecks, founder effects, and possibly other diversity-reducing impacts (Crossman & Harington, 1970; Skog et al., 2014).

Even in the most diverse population in the dataset (i.e., Interior Alaska; CHT), genome-wide genetic diversity is very low; per individual, a SNP is observed on average every 3.2 kb. In the eastern populations (e.g., St. Lawrence River; SLA), this number reduces to a SNP on average every 11.2 kb. This is remarkably low when compared to marine fishes including Atlantic herring *Clupea harengus* (1 variant per 0.3 kb; Martinez Barrio et al., 2016) and Atlantic cod *Gadus morhua* (1 variant per 0.5 kb; Star et al., 2011), as well as freshwater fishes such as rainbow trout *Oncorhynchus mykiss* (1 variant per 0.75 kb; Gao et al., 2018). Furthermore, northern pike were estimated to be on average 0.0089% - 0.032% different from the reference genome. For comparison, humans *Homo sapiens*, known to contain low levels of genetic variation, are four to 13-times more genetically diverse than the northern pike as assessed here (i.e., 0.11% - 0.13% difference from the reference; Auton et al., 2015). However, this low level of genetic diversity does not preclude evolutionary successful strategies; northern pike are renowned for their ability to thrive in a variety of environments and to expand into and colonize new areas (Luan et al., 2021).

In the region that separates high and low diversity North American northern pike, local effects may be further impacting diversity levels. The BC population Palmer Lake (PAL) is from a lake off the large Atlin Lake, which eventually flows into the Yukon River. The Palmer Lake population has lower genetic variation than the Yukon River population (HOO), even though these two sites are relatively proximal and may have at one time been directly connected. However, the Palmer Lake population may have lost variation due to inbreeding and drift. The apparent very low diversity of the Columbia River sample (CAS), and the lower diversity of Charlie Lake sample (CHA) relative to the Palmer Lake population may be due to technical artefacts, since both samples only had a single sample characterized from the population and underwent different sequencing technology to characterize their variation. Furthermore, the CAS individual was the source of the reference genome assembly, which would further reduce the characterized variation given that there would be no homozygous alternate genotypes observed. However, the Columbia River population (CAS) is in fact invasive from east of the NACD (Carim et al., 2019), and therefore it may have low diversity relative to BC populations west of the NACD. Additional samples would be needed from these two populations to accurately place them in the context of other populations genotyped here. Furthermore, introductions of northern pike have occurred in BC waters (Harvey, 2009), which could impact spatial trends in unexpected ways depending on the source population and size of the introduction. In conclusion, local impacts such as isolated populations or translocation history may explain differences from the expected level of diversity based on the broader regional location and proposed expansion history. The boundary region between elevated diversity and low diversity east of the NACD provides an interesting area of contrast that requires further characterization.

Although northern pike from east of the NACD had low genetic variation, in the present analysis each population was still consistently clustering with members of its own population with strong bootstrap support. Therefore, although previous studies have been unable to distinguish different populations of northern pike in eastern North America (Miller & Senanan, 2003; Senanan & Kapuscinski, 2000; Skog et al., 2014), in agreement with Ouellet-Cauchon et al. (2014) our results suggest separability of populations in the low diversity collections. The high degree of private alleles per population observed here is notable and may be explained by northern pike experiencing high genetic drift and expansion from a few individuals. These high levels however should be further investigated once more samples are included per population. Similar to distinct groupings observed here, although low nucleotide diversity was observed in eastern Russia, high genetic differentiation was observed in populations sharing a common origin, likely due to founder effects (Bachevskaja et al., 2019). Less differentiation may occur in central Europe. For example, in Germany, erosion of population substructure occurs alongside habitat degradation, which may be due to eutrophication and habitat loss leading to natural dispersal (Eschbach et al., 2021). The loss of differentiation also occurs by secondary contact of separate populations through human-mediated stocking, and diversity erosion was observed to be most severe in rivers or in modified water bodies (Eschbach et al., 2021). Given the well-supported clustering observed here across different populations in North America, population management may be feasible as reported for the congener *Esox masquinongy* (Rougemont et al., 2019). As an example, genetic markers have been used to determine the likely source of an introduced invasive population of northern pike in eastern Washington State, finding that it was likely a human-mediated transfer from disconnected populations in Idaho (Carim et al., 2022). To determine the ability to perform stock identification in the populations identified here would require further analysis with larger sample sizes to determine the expected power of genetic assignment tests (e.g., Moran & Anderson, 2019).

### Genetic diversity across the genome

Genetic diversity is not universally low across the northern pike genome, as certain regions are enriched for polymorphism (e.g., regions of chromosomes 5, 9, 11, and 24). These chromosomes have large regions of elevated diversity, which may indicate the presence of specific genes or high variability regions. Although a full inspection of the regions has not been conducted, some regions hold immune-related genes (e.g., major histocompatibility complex (MHC) region in chromosome 9). The MHC is known to be highly variable in vertebrates, and its variability is valuable for presentation of a diverse range of antigens for pathogen recognition (Piertney & Oliver, 2006; Unanue et al., 2016). Regions of elevated diversity in the genomes of otherwise low diversity species have been identified previously. For example, channel island foxes (*Urocyon littoralis*) have highest SNP density in olfactory receptor genes (Robinson et al., 2018). Humans, also known for low genetic diversity, have an enrichment of polymorphism around genes related to sensory perception and neurophysiology (Redon et al., 2006). Enrichment of polymorphism in certain areas is also observed in species with generally higher overall diversity; stickleback are known to have an enrichment of polymorphism around genes controlling bony plate number and morphology (Nelson et al., 2019). Collectively, these observations suggest that regions of high variation in the genome are associated with phenotypic variation that may be instrumental to the survival of the species. The maintenance of genetic variation may therefore be most crucial in genetic regions that affect the ability of a species to detect or respond to stimuli that are both fundamental to survival and that fluctuate in their habitat. The enriched polymorphic regions identified here in northern pike merit further investigation.

The observation of chromosome 24 having multiple separate regions of elevated diversity is in contrast with single regions as observed in chromosomes 5, 9, and 11. Importantly, in the populations that retain *amhby* in males, chromosome 24 contains the master sex determining gene (Pan et al., 2019). Differentiation of sex chromosomes in northern pike has previously been determined to be generally low (Pan et al., 2019). Lower levels of recombination between the X and Y chromosomes can occur, which may result in degeneration of Y due to the accumulation of mutations (Charlesworth, 1991). This can provide a benefit by sequestering sexually antagonistic alleles on sex chromosomes with reduced recombination (Charlesworth & Charlesworth, 2005; Mackay, 2001), but this is not expected in northern pike (Pan et al., 2019). In salmonids, for example, heterochiasmy occurs whereby a lack of recombination in males occurs throughout the genome, with crossovers primarily occurring at telomeric regions opposite to the centromere (Moen et al., 2004; Sakamoto et al., 2000; Sutherland et al., 2017). Heterochiasmy also occurs in humans (Broman et al., 1998). However, any heterochiasmy observed in northern pike appears to be low, as observed by Rondeau et al. (2014) that only a slight skew in recombination rates occurs between the sexes. The presence of multiple regions and generally elevated overall polymorphism enrichment in chromosome 24 merits further investigation, especially within populations that have lost the sex determining gene from this chromosome. With more populations and individuals characterized in the future, the precision of the location of these genomic regions or interpopulation variation that may exist will be valuable to be explored further.

### Sex determination and a patchwork of *amhby* presence in western North America

Range expansion, if impacted by founder effects or population bottlenecks, may not only result in a loss of neutral genetic variation but also in a loss of functional genetic variation. The master sex determining gene *amhby* is considered to have been lost during expansion into North America (Pan et al., 2021), which coincides with a loss of overall genetic diversity (Skog et al., 2014), and our results are concordant with this finding. Northern pike can undergo sex reversal, possibly influenced by environmental signals (Pan et al., 2019) as has been observed in other teleosts (Rajendiran et al., 2021), as well as through disruptions of other genes in the pathway including *amh receptor II* (Pan et al., 2023). Therefore, as described by Pan et al. (2021), the founding of a population comprised of *amhby*(-) individuals (i.e., no genetic males) followed by sex reversal could presumably result in the establishment of populations without the master sex determining gene.

In the present study, *amhby*(-) populations are largely what was observed in populations southeast to the Yukon Territory. However, this observation was not universal, as there were also detections of Alaskan *amhby*(-) males and Columbia River *amhby*(+) males. North American northern pike provides a valuable model to study sex determination evolution; as Pan et al. (2021) describe that the old genetic sex determining system is lost before a new system has taken its place, and environmental sex determination may play a transition state for these populations. The observations here point to a mosaic of *amhby* presence and presumed activity in males in the boundary region from Alaska through British Columbia. East of British Columbia, however, there were no observations of *amhby*(+) males. It is unclear how the Castlegar males retained *amhby*, as they may have retained the gene from the source population in Alaska, or they may be descendants of another lineage that still uses the *amhby* system. Further characterization of populations in this area may further elucidate the interaction between an environmentally mediated sex determination system and the potential for emergence of a new genetic system for sex determination (Pan et al., 2021).

The lack of the *amhby* gene in eastern populations does not coincide with a lack of males in eastern Canada. However, it is notable that upper St. Lawrence River populations have experienced sex ratio shifts from male to female dominance in recent decades (Farrell & Barry, 2012). An earlier attempt by Rondeau and co-workers (*unpublished*) to map the sex determining locus using five biparental crosses of St. Lawrence region northern pike where phenotypic sex was determined by histology resulted in three highly male-biased families (from 76-92% male) and two moderately male-biased families (from 56-65% male), and the inability to detect sex-linked markers was thought to be due to the strong sex bias combined with low polymorphism (Rondeau, *unpublished*). However, more recent analyses, as discussed above, suggest that this was due to the lack of a genetic sex determination system at play currently in these eastern populations. Additional analysis of these samples using the markers developed by Pan et al. (2021) also failed to detect *amhby*, and so the resequencing approach here was taken as reported in the present study.

### Genome assembly

A new individual was used for the latest version of the assembly due to decreasing quantity and quality of any remaining material used for earlier versions. Regardless, with the significant advances that occurred in sequencing and assembly, the latest assembly was superior in metrics and utility relative to earlier assemblies. In human medical, agricultural, or even now conservation and population genomic fields, there are an increasing number of assemblies for the same species, with researchers generating references specific to their population; this can have benefits in terms of alignment success and variant calling (Thorburn et al., 2023), but also needs to be considered in terms of detecting potential reference bias, in particular when populations are highly divergent (Bohling, 2020). In northern pike, there now exists both the North American female assembly from the Castlegar population of the Columbia River (presented here), as well as a high quality European male assembly (Pan et al., 2019) that was optimal for characterizing the sex determination region within the genome. Future additions of other assemblies from Asian or other lineages may eventually produce a pan-genome for the species that represents the diversity of this Holarctic species.

## Conclusions

In the present work we generate a long-read assembly with chromosome-length scaffolds, and use this resource to characterize population genetic trends at a genome-wide scale, as well as genomic diversity trends within and between chromosomes. By combining sex-specific analyses of whole-genome resequencing data with PCR-based assays for sex markers, we were able to identify a patchwork of populations with or without the male-based sex determination system *amhby*. This patchwork, so far including *amhby*(+) and *amhby*(-) males in Alaska and *amhby*(+) males in the Columbia River, points to a more complex landscape of the loss of this sex determining system that occurred from west to east in North America. The large number of private alleles and clear clustering of individuals within each population points to substantial drift in individual populations. Population distinctiveness would be expected to enable genetic stock identification of different populations, although this remains largely untested. The clear trends of decreased diversity (and loss of functional genetic sex determination gene) with distance from Alaska is in agreement with the theory that northern pike expanded outward from Alaska after the deglaciation of the last ice age, and that significant founder effects impacted populations in the recolonized range. Regional effects in population diversity were observed as well, and from evidence so far, the North American Continental Divide provides a marking point after which diversity is uniformly low. Additional samples from more southern regions including the Columbia River would be beneficial to understand these post-glacial trends.

## Acknowledgements

We are grateful to the following people and organizations for providing samples for this study: Charles O. Hayford State Fish Hatchery (Hackettstown, NJ, USA); the Alaska Department of Fish and Game; the Manitoba Fisheries Branch; Whiteshell fish Hatchery (West Hawk Lake, Manitoba); Jeremy Baxter of Mountain Water Research; and Marco Marrello of Terraquatic Resource Management. We thank Dr. Miya Qiaowei Pan and Dr. Yann Guiguen for collaboration and insights supporting the genetic sex analyses in this study. Thanks to Dr. Sergio Cortez Ghio of Ex Machina Biostats for discussions on identifying regions of polymorphism enrichment. This work was supported by NSERC (RGPIN/3888-2017) and the New York Environmental Protection Fund AM-10165 administered by the NYS Department of Environmental Conservation.

## Competing Interests

Ben Sutherland is affiliated with Sutherland Bioinformatics. The author has no competing financial interests to declare. The other authors declare that no competing interests exist.

## Data Availability

The reference genome v.4.0 is available through NCBI WGS accession SAXP01000000.1 and assembly accession GCA_004634155.1. Whole genome resequencing data is available through BioProject accession PRJNA512507 and SRA accessions SAMN10685075 – SAMN10685119.

The filtered VCF file with dataset SNPs from alignment to reference genome v4.0 is available on FigShare: https://doi.org/10.6084/m9.figshare.25230146

This work was supported by the following GitHub repositories:

General data analysis: https://github.com/bensutherland/ms_pike_popgen

Genome data analysis scripts: https://github.com/enormandeau/Scripts

## Author Contributions

Designed Research: Ben F Koop, Eric B Rondeau

Performed Research: Eric B Rondeau, Hollie A Johnson, Ben J G Sutherland, Joanne Whitehead, Cody A Despins, Brent E Gowen, Christopher M Whipps, John M Farrell, Brian J Collyard Analyzed Data: Hollie A Johnson, Eric B Rondeau, Ben J G Sutherland, David R Minkley, Jong S Leong, Joanne Whitehead

Wrote the Paper: Hollie A Johnson, Eric B Rondeau, Ben J G Sutherland

## Additional Files

**Figure S1.** Hi-C contact map as visualized in Juicebox v 1.8.8 for assembly v4.0 GCF_004634155.1.

**Figure S2.** Distribution of repeats along the assembly v4.0 GCF_004634155.1, displayed in bins of 100,000 bp along the chromosome, with counts of masked bases per bin.

**Table S1.** Primer names, sequences, and annealing temperatures for sex markers. SeqAMH1 and ConserveAMH1 are reported in Pan et al. (2021).

**Table S2.** RepeatMasker summary table for assembly version 4.0 (GCF_004634155.1). Description of custom repeat library generation and masking methods described in Rondeau et al, (2014).

**Additional File S1.** Counts and locus identifiers of heterozygous or homozygous alternate variants per individual for all individuals in the study.

**Additional File S2.** Sum of variants per window across the genome in 1 Mbp windows.

**Additional File S3.** Sex-specific SNPs identified from the regional analysis of the whole-genome sequence data, shown per genomic coordinate as a 0 (ref/ref) or 1 (ref/alt) for each sexed northern pike from Chatanika River.

## Notes

https://github.com/bensutherland/ms_pike_popgen

### Competing Interest Statement

The authors have declared no competing interest.

### Summary of Updates

This version of the paper provides a more detailed discussion of the insights gained from whole-genome resequencing data. The author list has been updated.

