## Supplementary figures and images for "Loss of genetic variation and sex determination system in North American northern pike characterized by whole-genome resequencing"

### Figure S1

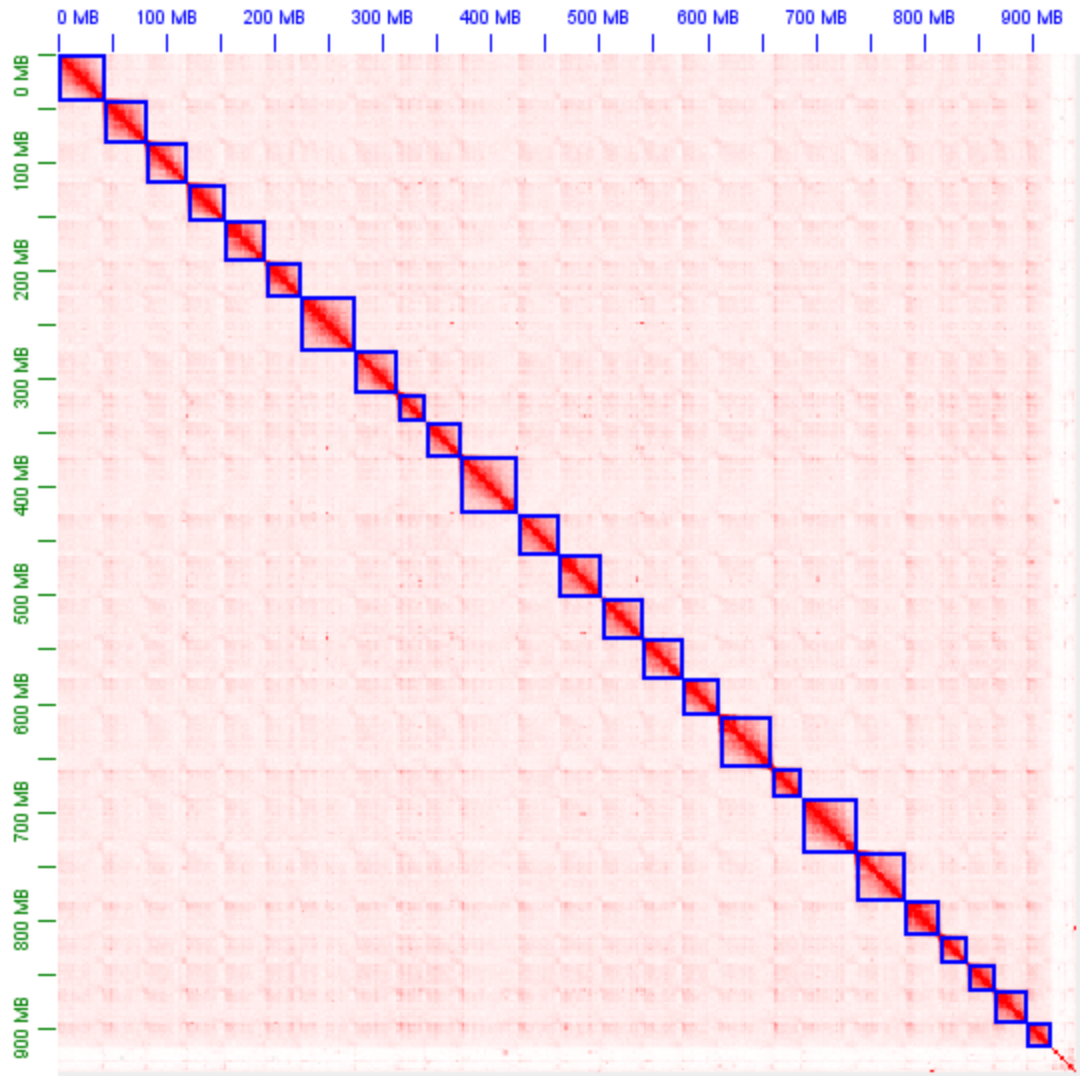

### Figure S2

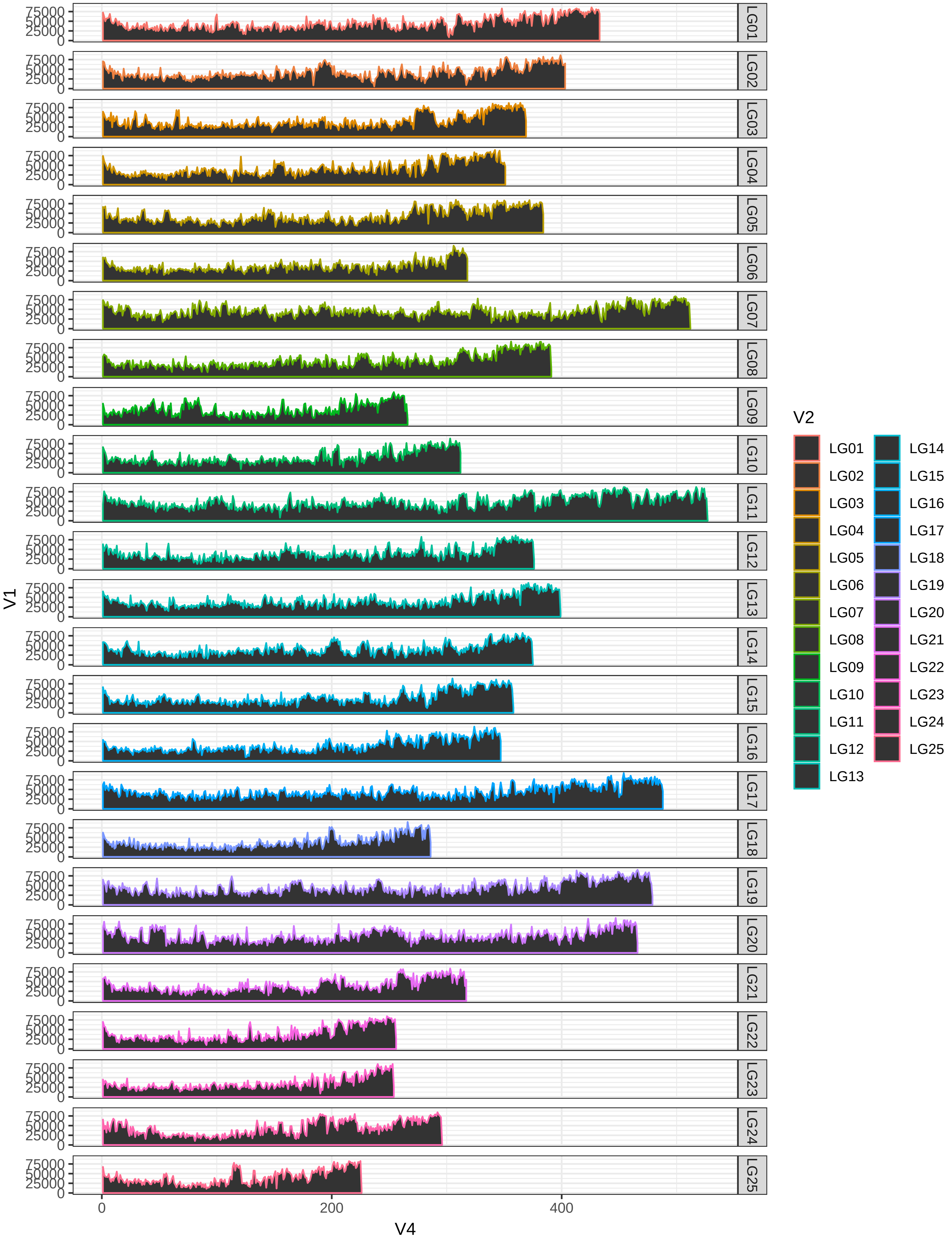
