## Supplementary material for "Loss of genetic variation and sex determination system in North American northern pike characterized by whole-genome resequencing": Table S1

**Table S1.** Primer names, sequences, and annealing temperatures for sex markers. SeqAMH1 and ConserveAMH1 are reported in Pan et al. (2021).

| **Primer Name** | **Sequence 5' → 3'** | **Annealing Temp. (˚C)** |
| --- | --- | --- |
| SeqAMH1Fw4 | CAACATGGTGGCAACTAAGTG | 52 |
| SeqAMH1Rev4 | GGTAATATTTGTGCCCTGTG |  |
| ConserveAMH1_F1 | GTTACTTTTTCTGCCTAGCGTGA | 54 |
| ConserveAMH1_R1 | CTATTACTAGTGTGGATAAGGCCG |  |
| 24.5 F | AATTACAGACCTCTACATGCT | 52 |
| 24.5 R | GATAGTCCCATAGATTGTAGA |  |
| 24.5 Probe | GCAAATGACGGCGCACTGTT |  |
